# Neural acceleration drives adaptations in working memory encoding speed

**DOI:** 10.64898/2026.06.17.732824

**Authors:** Joost de Jong, Mark Siertsema, Cemre Baykan, Elkan Akyürek

## Abstract

Humans and non-human animals adaptively boost their encoding speed when they expect limited sensory exposure time, so that they can capture essential information before it is gone. However, it is unclear how the brain implements this crucial adaptation. Using multivariate pattern analysis of human EEG data, we found that an accelerated neural code underlies adaptations in visual working memory encoding speed.

---

From watching a high-paced action movie to listening to a calming bedtime story, the rate at which information reaches the senses varies considerably. When organisms only have limited time to extract information from their surroundings, it is advantageous to accelerate neural encoding. Indeed, adaptations in encoding speed have been found across senses and species^1–3^, such as retinal light responses in mice^4,5^, perceptual decision-making in rats^6^ and humans^7–10^, and speech perception in humans^11^. Recently, we demonstrated that visual information is encoded in working memory more rapidly (i.e., encoding is accelerated) when human participants expect that information will be briefly available^12^.

However, it is not clear how the brain implements such a critical adaptation. Here, we contrasted two scenarios (see Figure 1): In the first scenario, neural encoding of information in visual working memory runs at the same speed (i.e., is not accelerated), but starts earlier, such that neural encoding is shifted in time throughout encoding (*early start scenario*). In the second scenario, neural encoding starts at the same time, but the underlying neural processes (transiently) run at a higher speed, gradually producing a temporal shift (*accelerated scenario*).

**Figure 1.**
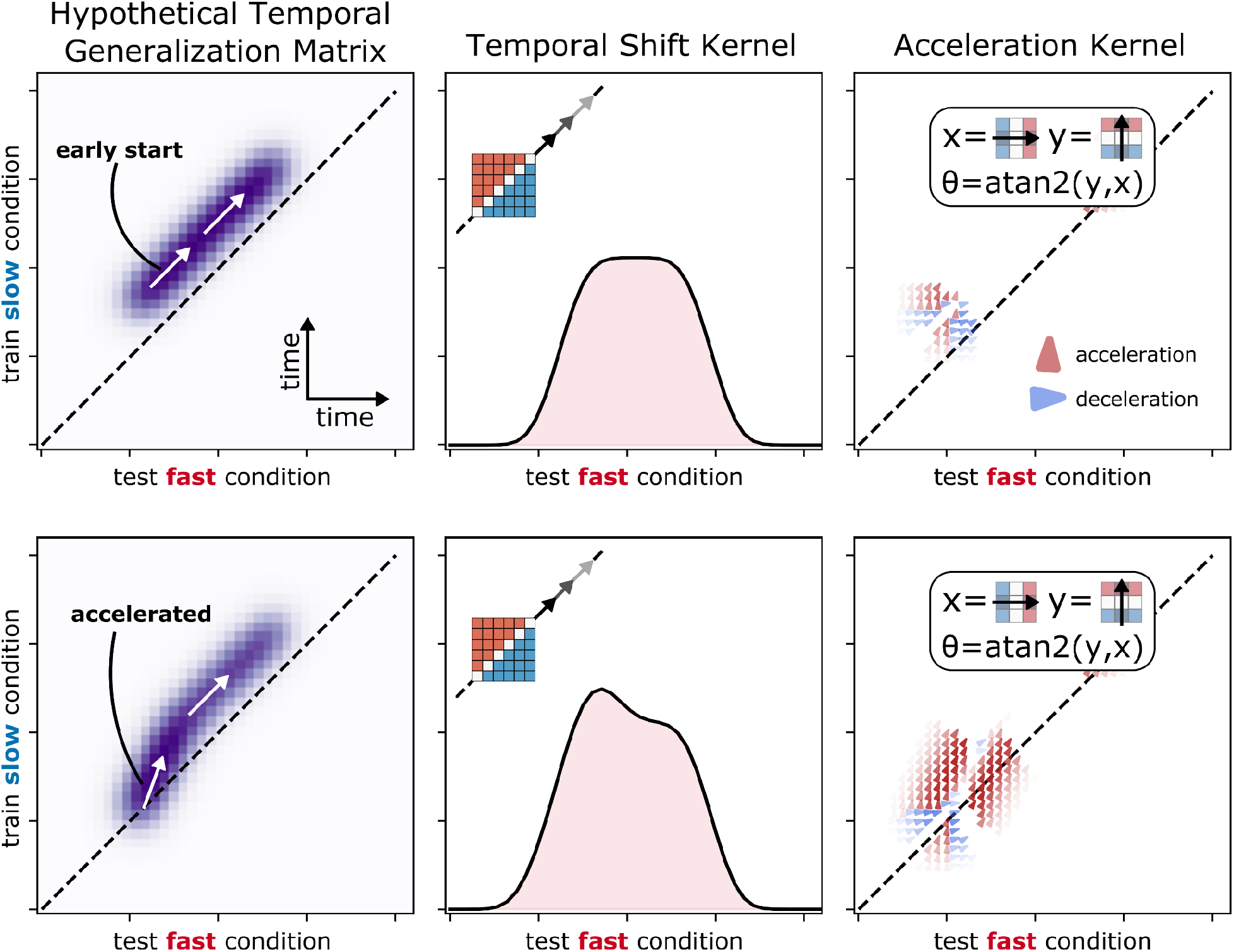
Two hypotheses (top and bottom row) concerning the neural mechanisms underlying adaptations in encoding speed, expressed in terms of temporal generalization matrices, where a decoder is trained at each timepoint and tested at all other timepoints. Left: synthetic temporal generalization matrices, with purple colors indicating high decoding scores, and the dashed line representing the diagonal where training time equals testing time. When the ‘fast’ condition is only shifted earlier in time, decoding is shifted from, but still parallel to the diagonal (top). When neural encoding is accelerated, decoding is initially on the diagonal, but because it proceeds at an angle with the diagonal, it eventually shifts above the diagonal (bottom). Middle: temporal shifts can be measured using a temporal shift kernel that slides across the diagonal. Because decoding centers above the diagonal, the shift score is positive. The shift kernel alone cannot distinguish between ‘temporal shift’ and ‘accelerated’ because it is too sensitive to overall decoding strength (the bulge on the left side of the bottom graph is visible, but does not necessarily signify a speed difference). Right: neural speed differences can be measured using an acceleration kernel that measures both the strength and angle of decoding gradients relative to decoding in the diagonal direction.

While both scenarios predict a temporal shift, they differ in whether that shift is there from the beginning (*early start scenario)* or whether the temporal shift is gained over the course of encoding through a neural speed difference (*accelerated scenario*). In this study, we use EEG, multivariate pattern analysis (MVPA^13^), and temporal generalization techniques^14^ to contrast both scenarios. To preview, our analyses suggest that when participants expect limited viewing time, neural encoding is accelerated.

To study adaptations in neural encoding speed, we asked participants to perform a visual working memory task^12^ while we measured neural activity using EEG. On each trial, a Gabor patch with random orientation was presented for 50, 200, or 400ms and subsequently masked by a random constellation of patches (see Supplementary Figure 1). After one second, participants reproduced the orientation of the presented Gabor patch using a scrolling wheel. The order of presentation times was counterbalanced, such that triplets of presentation times occurred equally often in each block (with 30 blocks × 30 trials). This allowed us to study how the temporal context, defined as the presentation time on previous trials, influenced encoding speed on the current trial (see Supplementary Figure 1).

If participants use the temporal context from previous trials to generate an expectation of how much time they have for encoding on the current trial, then encoding speed on the current trial should be faster when the previous visual targets were briefly exposed. This is exactly what we found behaviourally; short previous presentation times selectively increased encoding speed on the current trial (see Supplementary Figure 1). When the temporal context was ‘short’ compared to ‘long’, visual working memory performance on the current trial reached its asymptote quicker (*t*=3.97, *p*<0.001, BF_10_=193.19), without any changes in asymptote (*t*=0.42, *p*=0.68, BF_10_=0.14). These results replicate our earlier findings^12^ and suggest that humans can—over the course of just a few trials—implicitly boost visual working memory encoding speed.

But how does the brain accomplish speedups in visual working memory encoding? We used MVPA to decode the presented orientation from EEG activity in electrodes over visual areas (see Methods). Orientation was readily decodable from EEG and shows a remarkably dynamic neural code, with strong decoding primarily centered around the diagonal of the temporal generalization matrix (i.e., train time *equals* test time)^15,16^.

To test whether neural encoding is shifted in time, we leveraged the following logic^14^: In a temporal generalization matrix, on-diagonal decoding is expected if the train and test set share the same neural code and time course. But when neural processes in the test set are *shifted* earlier in time compared to the training set, we would expect orientation-specific neural patterns to surface earlier in the test set than in the train set. That is, neural decoding would exhibit off-diagonal decoding. A neural decoder was trained on long-context (i.e., long previous presentation times) trials, where visual working memory encoding speed was slow, and tested on short-context trials, where encoding speed was fast (see behavioural results). This cross-condition temporal generalization matrix exhibits stronger above-diagonal decoding if the neural encoding processes are shifted earlier in time in short-context trials compared to long-context trials (see Figure 1). To measure the degree of off-diagonal decoding, this matrix is then multiplied by a symmetric mask, ensuring that decoding scores outside of regions with significant decoding were discarded (decoding accuracy < 99th percentile; see Figure 2A). To quantify how much the decoding scores had shifted from the diagonal, we convolved the weighted matrix with a 35 × 35ms shift kernel along the diagonal (see Figure 2B), where positive values indicate above-diagonal decoding^17^. Positive shift scores indicate that the short-context condition has a time-shifted neural encoding, because the same neural code from some later training timepoint (long context) surfaces earlier in the testing trials (short context).

**Figure 2.**
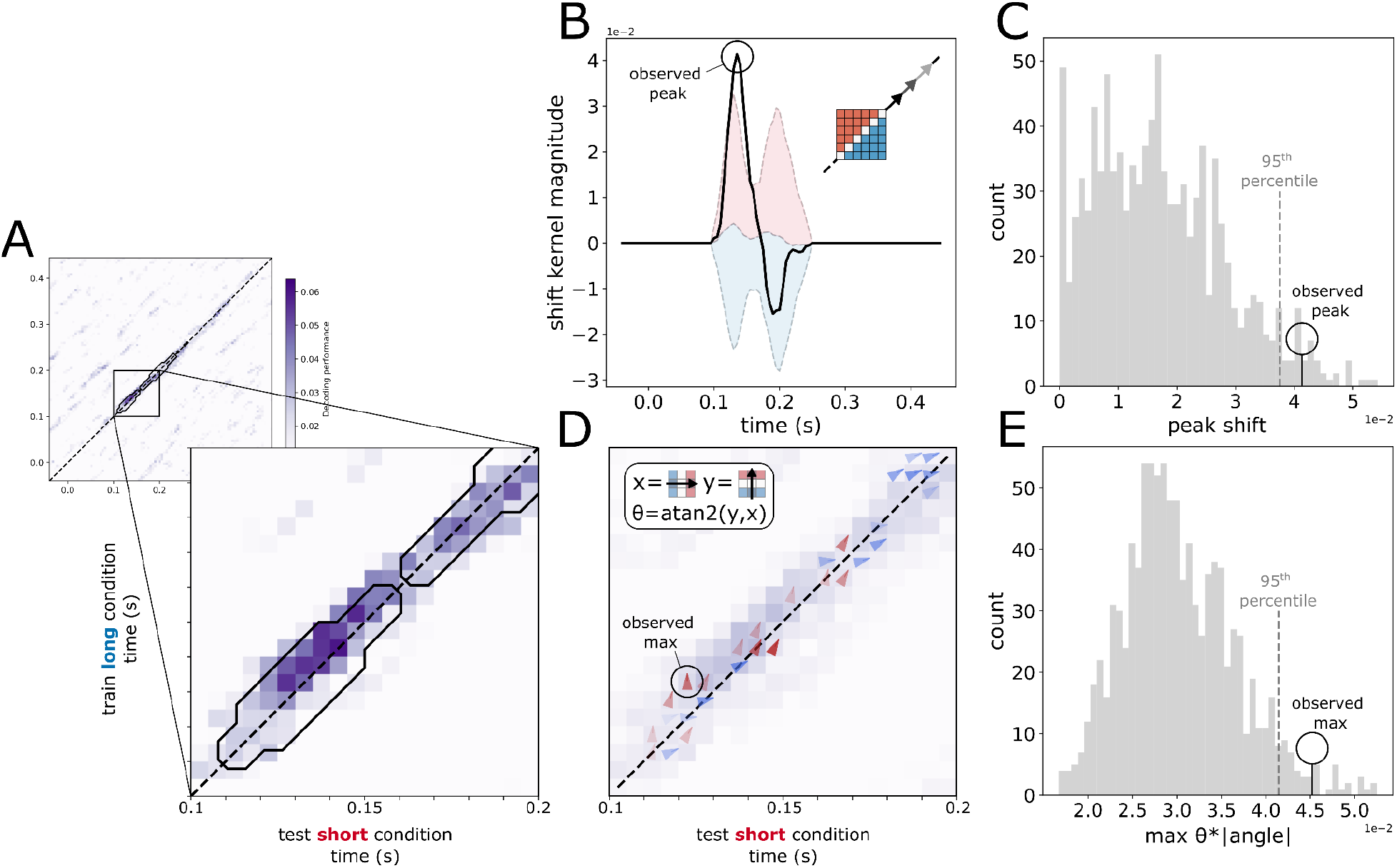
Neural encoding is accelerated when the temporal context is short. A: The cross-condition temporal generalization matrix, where the orientation decoder is trained on long-context trials and tested on short-context trials (only positive decoding scores are plotted). The inside of the black outline represents the mask used in our analyses. Above-diagonal decoding can be observed in the inset between 100 and 200ms. B: The temporal generalization matrix is convolved with a shift kernel, which computes the magnitude of above-diagonal decoding. Shaded regions represent permuted shift scores between the 5th and 95th percentile around the median. C: The observed peak is higher than the 95th percentile of peaks in the permutations. D: The temporal generalization matrix is convolved with an acceleration kernel, which measures the magnitude and angle of the decoding gradient (acceleration in red, deceleration in blue). E: The observed maximum angle is higher than the 95th percentile of maximum angles in the permutations.

We found that the aggregate cross-condition temporal generalization matrix (train = long context; test = short context) showed off-diagonal decoding between 100-200ms (Figure 2A). Indeed, the neural shift score peaked around 130 ms after stimulus presentation (Figure 2B). To test whether this peak in shift score was significantly different from what was expected by chance, we generated 1,000 permutations by swapping short and long context trials randomly between train and test sets, and found that the observed peak shift score was higher than the 95th percentile of permuted peak shift scores (P=0.028; permutation test; Figure 2C). Additional analyses suggest that this neural shift was not due to mask-related neural activity, and happened during visual presentation (P=0.002; permutation test; see Supplementary Figure 2). This suggests that, at around 130 ms after stimulus presentation, the neural pattern encoding stimulus orientation is significantly shifted in time.

Turning back to our main question, where does this temporal shift come from? Is neural encoding already shifted in time from the outset (*early start scenario*), or does neural encoding gain a temporal shift by running at a faster speed during encoding (*accelerated scenario*)? Here, we use the following logic^14^: When two neural processes run at different speeds, decoding in their cross-condition temporal generalization matrix should not only be located off the diagonal, but the *angle* of the decoding gradient should differ from the diagonal (i.e., different from 45 degrees; see Figure 1). In other words, for each single ‘decoding step’ in time in the training set, more than one ‘decoding step’ is taken in the test set. To measure neural speed differences (i.e., acceleration), we convolved the cross-condition temporal generalization matrix with an acceleration kernel that jointly measures the local strength and angle of the decoding gradient. Their product measures neural speed differences, where positive values reflect strong decoding rotated in the direction of an accelerated code for the short context (Figure 2D). To single out the decoding angles inside the region of highest decoding, we only considered angles inside the binary mask. We found that the maximum angle was bigger than the 95th percentile of the maximum of the permuted accelerations (P=0.021; permutation test; Figure 2E). This suggests that neural encoding gained a temporal advantage by being *accelerated* when humans expected limited viewing time.

To summarize, we found that adaptations in visual working memory encoding speed are driven by a neural acceleration. More specifically, the neural process that encodes information into visual working memory is accelerated when human participants expect limited viewing time. The acceleration of neural coding seems largely consistent with speedups found in, for example, retinal encoding of contrast and luminance^4,5^, and speech perception^11^, where neural patterns in response to visual or auditory information are sped up. Our findings also point to the striking functional role that time perception plays in adaptively tuning the speed of neural responses to the pace of our surroundings^18^.

It is not clear yet how adaptive acceleration is implemented at the neural level, but there are several candidate mechanisms, such as multiplicative gain on input and recurrent neural connections at the circuit-level^19–21^, neuromodulatory effects at the synaptic level^22,23^, or changes in the membrane time constant of individual neurons^24^.

Temporal speedups and scaling of neural responses have been found to account for a substantial portion of variability in neural activity^25^. Our results suggest that temporal neural variability is not entirely random, but instead reflects continuous adaptation to subjective temporal task demands. The temporal generalization methods we developed here enable straightforward testing of neural acceleration in a wide variety of domains, levels of abstraction, and species.

## Acknowledgments

We would like to thank Rebeka Kovács for assisting in data collection.

## Code and data availability

All data and analysis code are available on https://osf.io/r3e56.

## Online Methods

### Participants

32 first-year psychology students (22 females, 10 males; mean age 20.9 years) of the University of Groningen participated in exchange for partial course credits. Participants had normal or corrected-to-normal eyesight. The experiment was approved by the ethical committee of the Department of Psychology, and participants gave informed consent before the experiment started.

### Apparatus and Stimuli

The experiment took place in a sound-attenuated room with dimmed lights. Participants were seated approximately 60 cm from a computer monitor (19-inch CRT, 1280×1024 pixels, running at 100Hz) on which the stimuli were presented against a constant grey background (RGB = 128, 128, 128). To keep the distance to the screen constant and to reduce head movements, the participants had to place their chin on a chinrest and their forehead against a rubber band. The experiment was programmed in the free experiment builder OpenSesame^26^. Gabor patches with a Gaussian envelope (spatial frequency: 0.05 cycles/degree, standard deviation (std): 12 pixels, phase: 0) were presented at fixation. The orientation of the patch was chosen from a uniform distribution ranging from 0 to 180 degrees. Target patches were masked using 50 overlapping Gabor patches with their centres scattered in a 40×40 pixel square, and with their orientations randomised (spatial frequency: 0.05 cycles/degree, std: 10 pixels, phase: 0).

### Procedure

Before the experiment, participants signed a written consent form, after which the EEG cap was fitted. Participants were then seated in front of the monitor and were given verbal and written instructions. The experiment was the same delayed orientation estimation task as in our previous work^12^. The experiment consisted of 30 blocks with 30 trials each. An experimental trial started with the presentation of a fixation dot for 500 to 1000 ms (uniformly distributed), after which a Gabor patch was presented for 50, 200, or 400 ms. The order of presentation times of these patches was set via a de Bruijn sequence, implemented using software described by Aguirre and colleagues^27^. The de Bruijn sequence counterbalanced the order of elements of pre-set triples using the smallest sequence length. Here, triples of three presentation times were used (e.g., 50-50-200 ms), and all possible triplets (i.e., 3^3^ = 27 triplets) occurred with equal frequency within each block. This allowed for fitting counterbalanced encoding curves for each presentation time of the current trial n, using the presentation times from the previous two trials n-1 and n-2. The memory item was immediately followed by a pattern mask shown for 100 ms, suppressing retinal or cortical after-images, which partly controls the available encoding time for the memory item. After the mask, participants had to remember the orientation of the Gabor patch over the next 1000 ms, during which a blank screen was presented. Then, a Gabor patch that served as a probe was shown, which the participants could turn by scrolling the scroll wheel of the computer mouse. Participants were required to rotate this Gabor patch to match the orientation of the memorized Gabor patch, which they could confirm by clicking the left mouse button. Before the experimental trials, participants performed 12 practice trials, during which they received feedback on their responses after every trial. During the experimental trials, participants only received aggregate feedback after every block.

### EEG Acquisition and Pre-processing

EEG signals were recorded from 64 equidistant Ag/AgCl sintered electrodes on a CA-212 cap at a sampling rate of 1000 Hz. The data were recorded with *eego mylab* software, and an *eego* amplifier (EegoTMmylab | ANT Neuro). An electrode placed under the left eye was used as the electrooculography (EOG) channel, while an electrode on the left shoulder blade was used as the ground. The 5Z-electrode (Cz) on the top of the head was used as the online reference. We attempted to keep the impedance of all electrodes at or below 20 kΩ. Offline, the EEG signal was re-referenced to the average of both mastoid electrodes, bandpass filtered (Hamming window sinc FIR filter, 0.01 Hz high-pass, 30 Hz low-pass), and downsampled to 200 Hz using MNE-Python (version 1.6.1)^28^. Next, the data were epoched relative to the Gabor patch presentation onset at 0 ms (epoch from -300 to 1000 ms) and baseline corrected using the mean signal from -300 to 0 ms. EOG and electrocardiogram (ECG) artefacts were identified via independent component analysis (ICA). Here, the FastICA algorithm^29^ was applied to a copy of the EEG layout, which was only high-pass filtered at 1 Hz. Components were identified via visual inspection and comparison with the EOG signal, and subsequently removed from the epochs. The subsequent EEG analysis was performed using 17 posterior electrodes (10L, 9Z, 10R, 4LD, 9L, 8Z, 9R, 7RB, 5LC, 5LB, 8L, 7L, 7Z, 7R, 8R, 5RB, 6RB which correspond to 10-10 coordinates O1, Oz, O2, PO7, PO3, POz, PO4, PO8, P7, P5, P3, P1, Pz, P2, P4, P6, P8, respectively).

### Behavioural Analysis

The behavioural analysis was conducted in R (R Core Team, 2024, version 4.4.2) and is the same as in de Jong and colleagues^12^, with two exceptions. First, we noticed that some responses were likely outliers, which have a disproportionate effect on estimates of precision (σ^-1^), and therefore estimates of encoding speed and capacity. Therefore, for each participant, we excluded responses whose absolute error angle was more than Q75 + 5.5IQR (0.8% of all responses). We verified that this threshold minimized individual differences in encoding speed, and without this outlier exclusion, we still found a significant effect of temporal context on encoding speed (*t*=2.17, *p*=0.034, BF_10_=1.34). Second, some participants had such steep encoding curves that the nonlinear regression did not converge. To ensure convergence, we added implicit zero points for all participants at presentation time (t) = 1ms. Again, we still found a significant effect of temporal context on encoding speed without adding this implicit zero point (*t*=2.79, *p*=0.007, BF_10_=6.13).

To calculate the error angles, the angle of the memory item was subtracted from the response angle and multiplied by two (because Gabor patches have a rotational symmetry of 180 degrees, instead of 360 degrees), resulting in an error angle range of -180 to 180 degrees. Memory precision was then taken as the inverse of the circular standard deviation (σ^-1^) of the error angles per presentation time (t). To correct for random guessing, the memory precision that would be expected for a certain sample size N was subtracted from the sample. This was done by taking the inverse of the mean of 1,000 circular standard deviation estimates from circular uniform distributions with sample size N.

In order to contrast trials in which participants expected short versus long presentation time, we defined the ‘short’ temporal context (i.e., n-1 and n-2 presentation time were 50-50, 50-200, or 200-50ms) and ‘long’ temporal context (i.e., n-1 and n-2 presentation time were 400-400, 400-200, or 200-400ms; see Supplementary Figure 1). This also allowed us to use one condition as a training set and the other as a testing set in the neural decoding analyses, which is essential to measuring neural shifts and accelerations.

To quantify encoding capacity and speed in working memory, exponential encoding curves^30^ were fitted per participant, per temporal context, using the two parameters of maximum capacity (c) and encoding speed (*τ*^-1^):

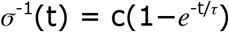

These parameters map onto the asymptotic memory precision for long presentation times (this asymptote being the maximum capacity *c*), and encoding speed (the speed at which precision reaches that asymptote; *τ*^-1^). The impact of maximum capacity (c) becomes most noticeable for longer presentation times as stimuli tend to be remembered more effectively after being presented for a longer duration. On the other hand, the effects on encoding speed are most prominent for shorter presentation times, as two encoding curves may eventually reach the same asymptote but at different rates. For instance, a fast encoding curve achieves a high level of precision early on; in contrast to a slow encoding curve, which reaches the same asymptote at a later timepoint. When fitting the encoding curves, these distinct behavioural characteristics of capacity and encoding speed allow for a reliable differentiation between the two.

In order to fit the encoding curves, we used the nls.multstart R-package v.1.3.0^31^, which allows for initializing a range of values for each parameter when fitting the encoding curve and selecting the pair of parameters that resulted in the lowest Akaike information criterion (AIC). Because the number of parameters was constant, the lowest AIC score translated into the highest likelihood. To assess the changes in capacity and encoding, the parameters were normalised by dividing the estimates by the mean estimate per participant. Finally, a linear regression model was used to evaluate the effect of information rate on the normalised encoding speed. Here, the null model predicts normalised encoding speed using only the normalised capacity, while the alternative model also uses the manipulation of temporal context (i.e., previous presentation times). Subsequently, approximated Bayes Factors^32^ were used to quantify the evidence for a modulation of the encoding speed as a function of the temporal context (short / long): BF_10_= exp(BIC_0_-BIC_1_) / 2. Here, BIC_0_ is the Bayesian Information Criterion score for the null model (with only normalized capacity as a predictor), while BIC_1_ also includes the temporal context (short or long). High BF_10_ values indicate evidence against the null and thus imply the presence of an effect of the expected presentation time on encoding speed.

### Multivariate Pattern Analysis

We performed Multivariate pattern analysis (MVPA) to determine whether the EEG signal from the posterior electrodes contained information on the to-be-remembered orientation of the Gabor patch. We used regularised circular regression to decode orientation, using the algorithm developed by Presnell and colleagues^33^ and implemented in Python by Alex Williams (https://gist.github.com/ahwillia/2941fb64e8bfe999c66291257601dfb4; MIT license). We used a rolling window of EEG voltages at posterior channels, centered and normalized around each timepoint, spanning -25 to 25ms^34^. Decoding accuracy scores varied between -1 and 1, where positive values indicate that the circular regression model fits well, and therefore that we can accurately decode orientation from EEG activity at posterior electrodes.

### Generalisation across time and conditions

To quantify the neural code for orientation over time, we trained our orientation decoder (using all presentation times) on all timepoints using a subset of trials, and for each trained timepoint, we tested on all other timepoints in the held-out testing set. We used MNE-python for generalization across conditions and time^28^. Such a temporal generalisation analysis^14^ results in a cross-temporal matrix of training time × testing time. To quantify neural shifts and accelerations, we generated a cross-condition temporal generalization matrix by training our orientation decoder on trials in which the temporal context was long, and testing on trials in which the temporal context was short.

### Measuring temporal neural shifts

Before computing the neural shift score, we weighed the cross-condition temporal generalization matrix by a symmetric, binary mask, which ensured that the shift score is mainly sensitive to regions in which significant orientation decoding is observed. For each participant, using all data, we computed the mean of five temporal generalization matrices through five cross-validations. Then, the median temporal generalization matrix over all participants was computed. To make the mask symmetric, we computed the mean of the top and bottom triangles of the matrices and mirrored the resulting triangle, making it symmetric. Then, any decoding that fell below the 99th percentile of decoding scores in the symmetric mask were set to zero, and any scores above the 99th percentile were set to one.

To quantify the degree of off-diagonal decoding, we adapted the neural shift score method as developed by Wolff & Akyürek^17^. This method computes the signed difference between decoding scores above and below the diagonal in a temporal generalization matrix. This is similar to convolving a Prewitt kernel of size k (which is used in signal processing to detect intensity gradients in images) along the diagonal of our matrix, where the values in the kernel are 0 along the diagonal, 1/k in the upper triangle, and -1/k in the lower triangle. The output of the kernel is positive when there is a positive decoding gradient going from the lower to the upper triangle. In other words, when kernel output is positive, decoding is clustered above the diagonal, which suggests a relative neural shift in time. The size of the kernel in all analyses was 7 × 7 (35ms × 35ms). Statistical inference was performed using permutation testing, because 1) the null-distribution for shift scores is not known, and 2) it allows for appropriate multiple comparisons correction. We randomly shuffled the short and long trials around between train and test sets 1,000 times. For each permutation, we computed the temporal generalization matrix, multiplied it with the symmetric mask, and computed the shift score, generating a null-distribution of 1,000 time courses of the shift score. We tested whether the observed peak shift was higher than the 95th percentile of the peaks in the permutations^34^. This permutation test amounts to a one-sided test, since we hypothesized that the neural code in the test (short condition) would be shifted earlier in time (i.e., have a positive value).

### Measuring accelerated neural codes

To quantify neural speed differences, we measured the strength and orientation of the local gradient of decoding in the cross-condition temporal generalization matrix by convolving it with a horizontal (x) and vertical (y) Sobel kernel. The angle of the local gradient can then be computed by taking the atan2 of vertical and horizontal gradients. The angle of the local gradient was then transformed such that 1) 0° corresponds to the diagonal direction and positive angles correspond to an accelerated neural code for the test set, and 2) angles larger than 45° or smaller than -45° are discarded, because they correspond to time-reversed neural codes. The resulting angles were multiplied by the strength of the gradient, which is 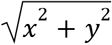. To ensure that only acceleration scores around the region of highest decoding were considered, we multiplied acceleration scores by the binary mask (see previous section). We used the same permutations as for the neural shift scores, and checked whether the observed maximum acceleration score was higher than the 95th percentile of the maximum acceleration score in the permutations. This permutation test amounts to a one-sided test, since we hypothesized that the neural code in the test set (short condition) would speed up (i.e., have a positive value).

## Supplementary figures

**Supplementary Figure 1.**
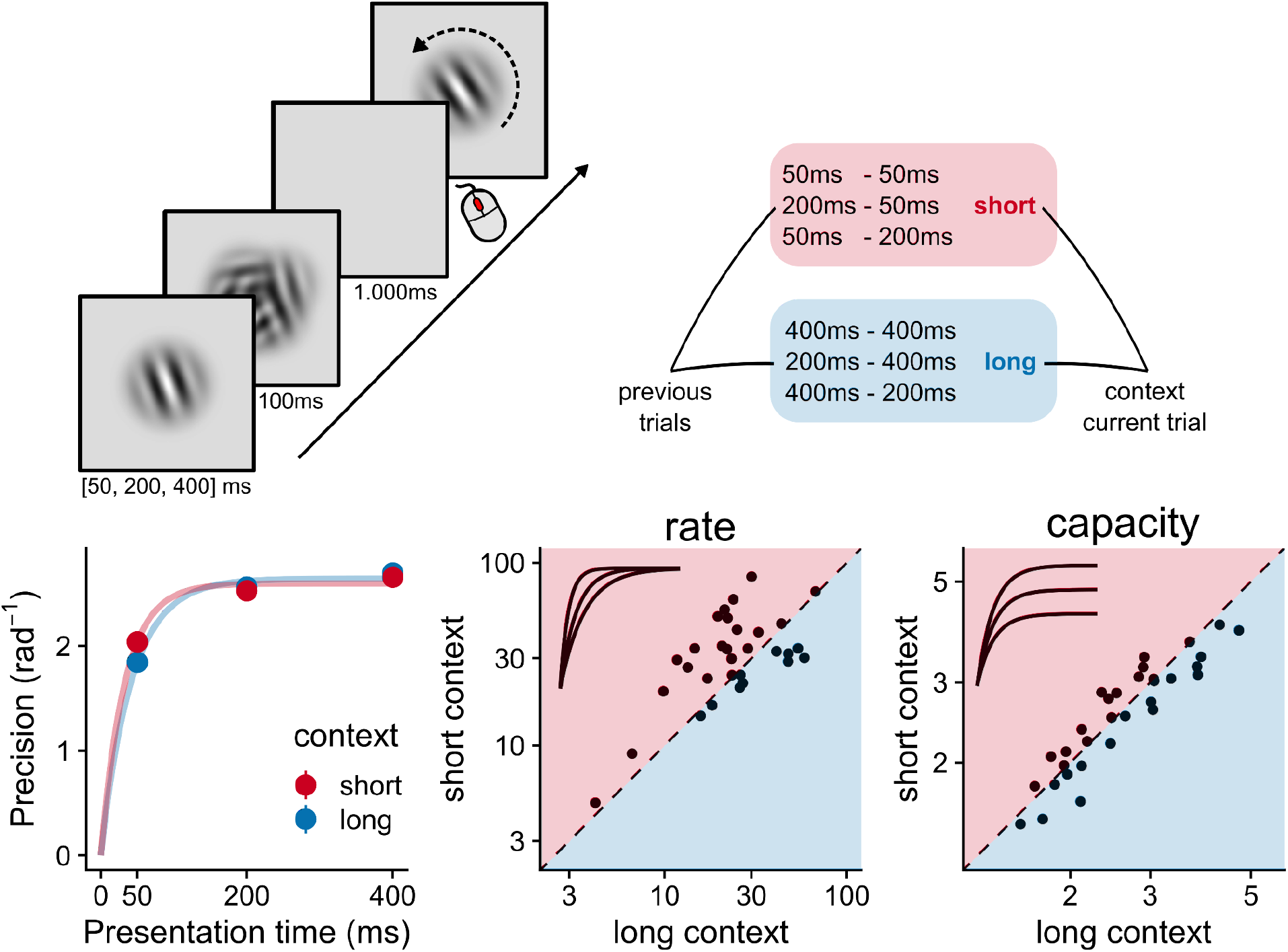
Humans adapt visual working memory encoding speed to expected viewing time. Top left: On each trial, participants were shown an oriented Gabor patch for 50, 200, or 400ms, which was subsequently masked. After one second, they needed to rotate a second Gabor patch using a scrolling wheel to match the memorized orientation. Top right: Previous trials had either short or long presentation times, setting the temporal context for the current trial. Bottom left: precision of the reproduced orientation was captured by an exponential encoding curve with two parameters: Rate and capacity. Bottom middle: The rate at which visual working memory performance (precision) reached its asymptote was higher in the ‘short’ condition than in the ‘long’ condition. Bottom right: The capacity (i.e., the asymptote) did not differ between conditions.

### Neural acceleration happens during visual presentation

Our previous behavioural experiments on adaptive encoding speed in visual working memory suggest that, in addition to a speedup during encoding, post-encoding processes such as visual masking may play a role^12^. However, it is difficult to disentangle these factors based on visual performance alone, which is necessarily a low-dimensional summary of a finely timed chain of neural processes. Therefore, we checked trials in which the mask could not have any influence on neural activity during peak decoding, namely when the stimulus was presented for 400ms. Using only a third of the total number of trials for this analysis, we found a highly reliable temporal shift peaking at 190ms (P=0.002; permutation test), with significant shifts observed from 175ms to 210ms (Supplementary Figure 2). However, we did not find a reliable acceleration during neural encoding (P=0.079), possibly due to reduced power (this analysis used a third of the trials used in the main analyses). The more pronounced neural shift is likely due to reduced influence of mask-related neural responses and occurs much earlier than mask onset at 400ms. Hence, our observed neural shifts are largely due to faster encoding of visual information, independent of ensuing visual responses or eye movements^36^.

**Supplementary Figure 2.**
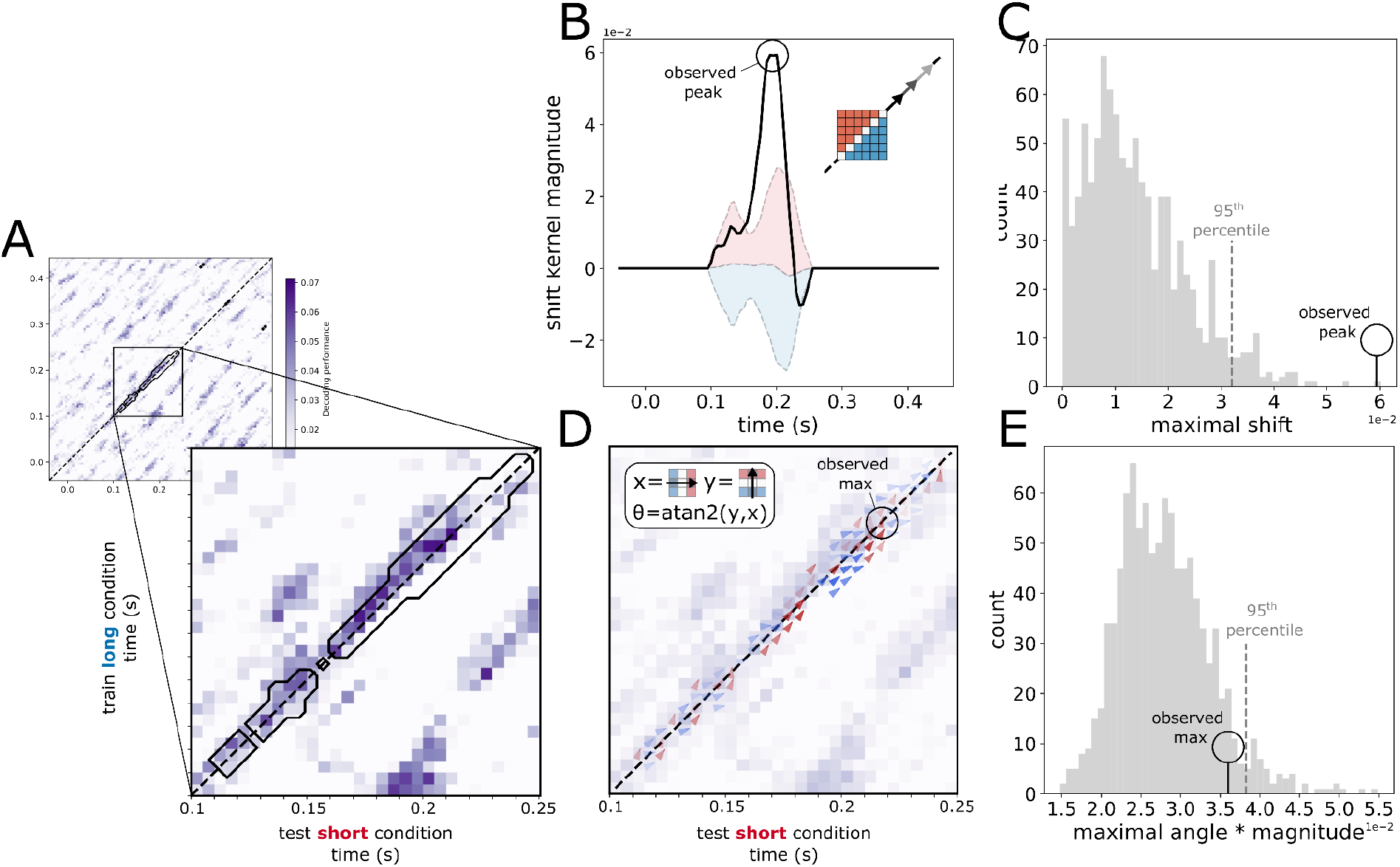
Neural encoding is accelerated when the temporal context is short, when the stimulus is presented for 400ms. A: The cross-condition temporal generalization matrix, where the orientation decoder is trained on long-context trials and tested on short-context trials (only positive decoding scores are plotted). The black outline represents the mask used in our analyses. Above-diagonal decoding can be observed in the inset between 100 and 200ms. B: The temporal generalization matrix is convolved with a shift kernel, which computes the magnitude of above-diagonal decoding. Shaded regions represent permuted shift scores between the 5th and 95th percentile around the median. C: The observed peak is higher than the 95th percentile of peaks in the permutations. D: The temporal generalization matrix is convolved with an acceleration kernel, which measures the magnitude and angle of the decoding gradient (acceleration in red, deceleration in blue). E: The observed maximum angle is higher than the 95th percentile of maximum angles in the permutations.

## Notes

https://osf.io/r3e56

### Competing Interest Statement

The authors have declared no competing interest.

